# GABAergic inhibition differentially gates recruitment of dentate gyrus interneurons by lateral entorhinal cortex inputs

**DOI:** 10.64898/2026.03.11.710817

**Authors:** Johannes Köhler, Marlene Bartos, Claudio Elgueta

**Affiliations:** Institute for Physiology I, University of Freiburg, Medical Faculty, Freiburg 79104, Germany; Institute for Anesthesiology and Perioperative Medicine, University Hospital of Zurich, Zurich 8091, Switzerland

**Keywords:** Dentate gyrus, interneurons, pattern separation, GABAergic inhibition, feedforward inhibition, feedback inhibition, parvalbumin interneurons, optogenetics, lateral entorhinal cortex

## Abstract

By keeping its activity low even when strongly stimulated, the dentate gyrus (DG) serves as the main entry point to the hippocampal circuit and helps separate different patterns of incoming information. However, the mechanisms supporting this low-level activity remain unclear. Here, we used *in vitro* patch-clamp recordings, two-photon calcium imaging, and optogenetics in hippocampal slices from adult mice to understand how local GABAergic interneurons control lateral entorhinal cortex (LEC) input-mediated drive in the DG. Under control conditions, LEC inputs rarely elicited granule cell (GC) firing because GABA_A_ receptor-mediated inhibition strongly restrained GC excitability. Whole-cell patch clamp recordings of DG interneurons revealed that LEC activation prevalently recruited fast-spiking parvalbumin-expressing and molecular layer interneurons, while most dendrite-targeting interneuron types responded only weakly to LEC input and fired action potentials only after feedback excitation from GCs was enhanced by pharmacological block of GABA_A_-receptors. Optogenetic inhibition of defined interneuron populations showed that silencing dendrite-targeting interneurons caused a larger increase in GC population spiking than silencing perisomatic-targeting interneurons, suggesting that both molecular layer and feedback-recruited interneurons have a prevalent role in controlling DG recruitment after an increase in LEC drive. In summary, our data indicate that GABAergic inhibition engages distinct DG interneuron types in feedforward and feedback circuits to tightly gate entorhinal inputs. By maintaining sparse GC firing, this dynamic inhibitory gating supports the role of the dentate gyrus in the hippocampal pattern-separation process.

## Introduction

Sparse neuronal networks are highly efficient at storing large numbers of distinct representations by performing pattern separation (Treves and Rolls, 1994; Rolls, 2013). The hippocampal dentate gyrus (DG) is a prominent example of this concept, as granule cell (GC) activity during spatial navigation is markedly lower compared to other downstream hippocampal and cortical areas (Pernía-Andrade and Jonas, 2014; Diamantaki *et al*., 2016; Senzai and Buzsáki, 2017; Hainmueller and Bartos, 2018). This sparse activity, in combination with absent synaptic GC-GC connections, enables the DG to support the emergence of orthogonal spatial representations, even when environmental changes are subtle and below the threshold for behavioral output (Leutgeb *et al*., 2007; McHugh *et al*., 2007; Allegra *et al*., 2020).

The DG receives the majority of its excitatory inputs via the perforant path (PP), which originates from the medial and lateral entorhinal cortex (MEC and LEC, respectively) and projects to the middle and outer molecular layers (Witter, 2007). The MEC conveys allocentric spatial information while the LEC provides signals related to objects, cues, or defined events, thereby functionally segregating the inputs entering the DG (Hargreaves *et al*., 2005; Deshmukh and Knierim, 2011; Wang *et al*., 2018). Although the hippocampus can transform these dynamic EC inputs into stable, sparse spatial maps (Cholvin *et al*., 2021), the precise mechanisms by which the DG integrates these highly active inputs while maintaining a sparse coding scheme remain unclear.

Various cellular and circuit properties may underlie the sparse activity of GCs and the selective recruitment of cells within an engram encoding a specific environment (Liu *et al*., 2012). A hyperpolarized resting membrane potential (Pernía-Andrade and Jonas, 2014), combined with strong dendritic attenuation and low dendritic excitability (Schmidt-Hieber *et al*., 2007; Krueppel *et al*., 2011), is a factor that likely contributes to the sparse recruitment of GCs following EC activation (Zhang *et al*., 2020). Additionally, GCs are densely innervated by a diverse population of GABAergic interneurons (INs), providing both feedforward and feedback inhibition (Bartos *et al*., 2001; Bartos and Elgueta, 2012; Hosp *et al*., 2014; Savanthrapadian *et al*., 2014; Elgueta and Bartos, 2019). Fast-spiking parvalbumin-expressing interneurons (PV-INs), including basket cells (BCs) and axo-axonic cells (AACs, Bartos and Elgueta, 2012; Elgueta, Köhler and Bartos, 2015), mediate perisomatic inhibition, while somatostatin- (SOM, Savanthrapadian *et al*., 2014; Yuan *et al*., 2017), cholecystokinin- (CCK, Hefft and Jonas, 2005), and neurogliaform INs (Armstrong *et al*., 2011) target distinct domains of GC dendrites and provide dendritic inhibition. Most DG INs extend their dendrites to both the molecular layer—enabling feedforward activation by EC inputs—and the hilus, where they can be activated by collateral projections from GC axons or mossy cells, forming powerful feedback inhibition loops (Scharfman, 2016).

Interestingly, while PV-INs strongly influence the recruitment and synchronization of GCs during fast network activity patterns and provide lateral inhibition (Bartos *et al*., 2007; Strüber *et al*., 2015, 2017), they have minimal impact on the number of GCs active in representing a spatial location, as indicated by cFos labeling (Stefanelli *et al*., 2016) and *in vivo* recordings (Hainmueller *et al*., 2024a). Conversely, SOM-Ins, which target the distal portions of GC dendrites, effectively regulate the sparsity of DG assemblies and are essential for spatial memory discrimination (Stefanelli *et al*., 2016; Morales *et al*., 2021; Hainmueller *et al*., 2024a). This highlights not only the critical role of inhibitory INs in shaping DG activity but also underscores their significance in determining the size of active DG assemblies by inhibiting LEC distal inputs.

Here, we combined in vitro whole-cell patch-clamp recordings, two-photon Ca²⁺ imaging, pharmacological interventions, and optogenetic manipulations to investigate how LEC inputs are processed within the DG circuit. Our findings demonstrate that GABAergic inhibition not only effectively controls GC activity following LEC recruitment but also differentially modulates the activity of distinct IN subtypes within the DG.

## Results

### GABA_A_R-mediated inhibition controls the sparsity of GC activity after LEC activation

In vivo whole-cell recordings in awake, running mice have shown that although GCs display low levels of action potential (AP) firing, they continuously receive strong gamma-(30-100 Hz) and theta- (3-12 Hz) modulated synaptic inputs (Pernía-Andrade and Jonas, 2014; Zhang *et al*., 2020). However, the role of GABAergic inhibition in maintaining these low levels of GC activity remains unclear. To investigate the influence of GABA_A_ receptor (GABA_A_R)-mediated inhibition on GC activity, we stimulated LEC projections within the lateral perforant path (LPP) by placing an extracellular electrode in the outer molecular layer while recording evoked Ca^2+^ responses in GCaMP-expressing GCs, using a 2-photon scanning microscope (**Fig. 1a**). Under control conditions, gamma-modulated stimulation patterns (5 pulses at 30 Hz) failed to elicit detectable Ca^2+^ responses in GCs at low intensities (0.03 ± 0.01 and 0.04 ± 0.01 dF/F at 25 and 50 V stimulation intensity). At higher intensities, stimulation induced a mild increase in fluorescence, suggesting that some GCs reached AP firing thresholds (0.09 ± 0.01 and 0.16 ± 0.01 dF/F at 75 and 100 V stimulation intensities, **Fig. 1b**). Blocking GABA_A_Rs with gabazine significantly increased both the number of responsive cells and the amplitude of Ca²⁺ signals across all tested stimulation intensities (p < 0.001, **Fig. 1b**), consistent with a substantial increase in AP discharges from GCs. These results indicate that recruitment of the DG circuitry by LEC inputs is tightly controlled by GABA_A_R-mediated inhibition provided by INs. In subsequent experiments, we will investigate how the diverse population of DG INs provides inhibitory modulation of GCs and how this modulation is itself regulated by GABA_A_R-mediated inhibition.

**Figure 1:**
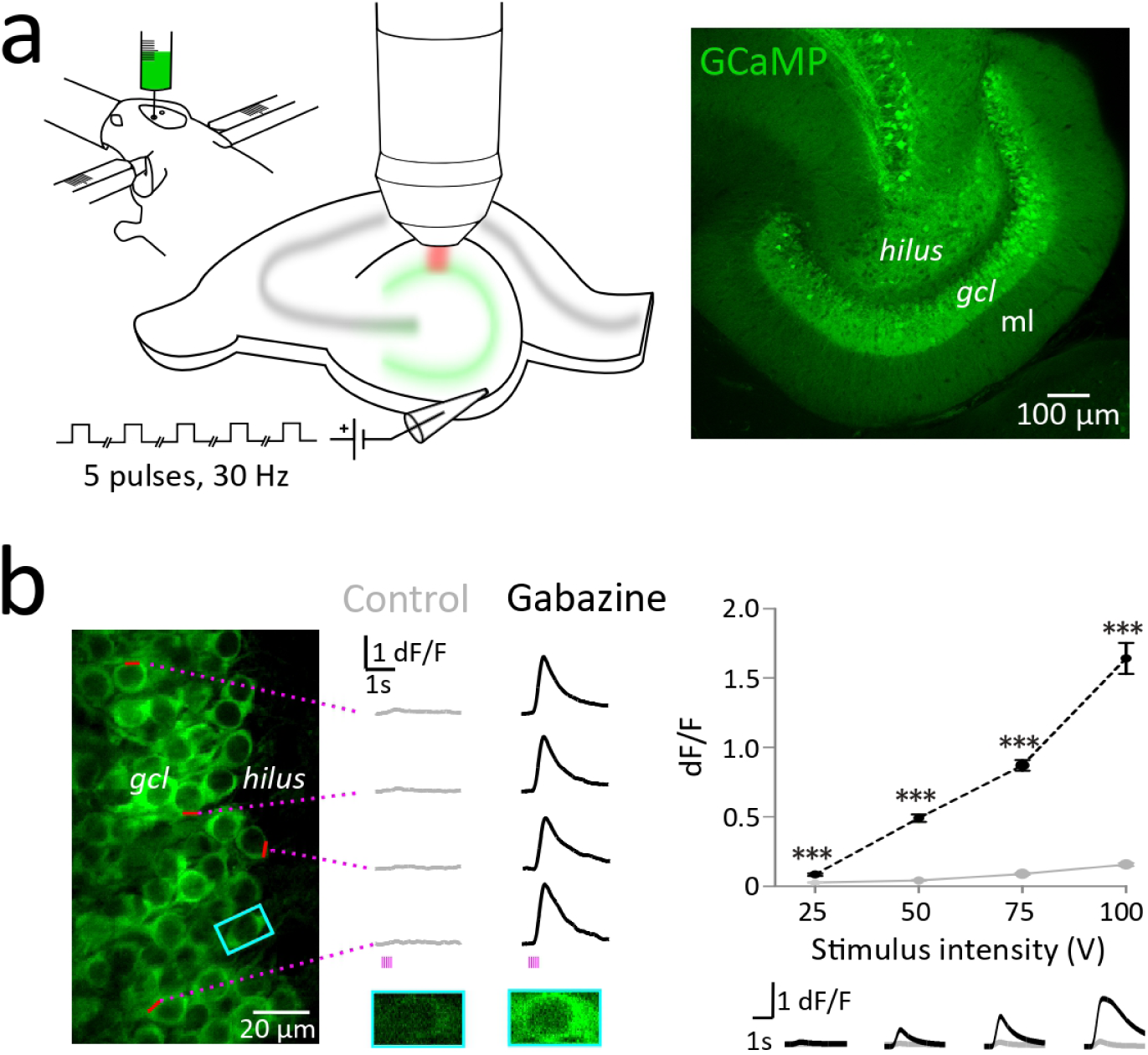
GABA_A_R-mediated inhibition controls the recruitment of DG GCs by LEC inputs. **(a)** *Left:* Schematic of the experiment. Ventral hippocampal slices were prepared from adult mice after stereotactically injecting adeno-associated viruses (rAAVs) encoding GCaMP5 into the ventral DG of adult mice. LEC fibers were activated using extracellular gamma-modulated stimulation (5 pulses at 30 Hz) in the outer molecular layer. *Right*, maximum intensity projection of confocal sections showing expression of GCaMP5 in the DG. The hilus, granule cell layer (gcl), and molecular layer (ml) can be seen. **(b)** Gamma-modulated activation of LEC inputs produced negligible increases in Ca^2+^ signals (dF/F) under control conditions (grey). Perfusion with Gabazine (SR-95531, 5 µM) resulted in a substantial increase in the Ca²⁺ responses (black) to LEC stimulation across all tested intensities. n=261 cells, 5 slices, 2 animals. *** p < 0.001. Two-way ANOVA followed by Holm-Sidak comparisons.

### DG Interneuron morphological and physiological characterization

GABAergic INs in the DG comprise a heterogeneous population with distinct morphologies, neurochemical profiles, and physiological properties (Hosp *et al*., 2014). To investigate how different types of INs may provide inhibitory modulation of GCs, we recorded INs in acute slices from adult mice by targeting neurons located at either the hilar-granule cell layer (*gcl*) border or in the molecular region of the DG. Post-hoc biocytin labeling after recordings allowed us to identify and classify INs based on their morphological properties, including their axonal projections. We detected six main IN types in our sample. BCs, with axons that project to the *gcl* and surround GC somas (**Fig. 2b**); AACs, whose axons cover the lower part of the *gcl* and form dense radially oriented cassettes (**Fig. 2c**); molecular layer cells (MLs), whose soma, dendrites, and axonal projections are located exclusively in the molecular layer (**Fig. 2d**); total molecular layer cells (TMLs), with axonal projections broadly spanning the molecular layer (**Fig. 2e**); and hilar-commissural PP-associated cells (HICAP), whose axons specifically cover the inner molecular layer (**Fig. 2f**). Additionally, we recorded SOM interneurons (SOMIs) using a reporter line that expresses the fluorophore tdTomato specifically in these cells (see **Materials and Methods**). First, we analyzed the physiological properties of cells with clearly distinguishable morphology (**Fig. 2**, **Table 1**). GCs were electrophysiologically identified based on their hyperpolarized resting membrane potential (RMP) and low firing rates (**Fig. 2g, h**, **Table 1**), as well as their higher accommodation index during intracellularly evoked firing patterns (**Fig. 2i**, **Table 1**). In contrast, all INs—except for ML cells—showed a significantly more depolarized RMP (**Fig. 2g**). Although no single physiological parameter was sufficient to distinguish individual IN morphological types, BC and AAC cells displayed a higher frequency discharge pattern (FS-INs), narrower spikes, and lower membrane resistance compared to TML and HICAP, which showed a regular-spiking pattern (RS-INs), and compared to ML cells. SOMIs exhibited a broader distribution of their physiological parameters, supporting previous findings indicating that they constitute a heterogeneous cell population (Yuan *et al*., 2017). On average, SOMIs displayed a phenotype that lies between FS- and RS-INs. Because physiological properties do not reliably separate IN types, this study relies primarily on morphological criteria for IN classification, unless otherwise indicated.

**Figure 2:**
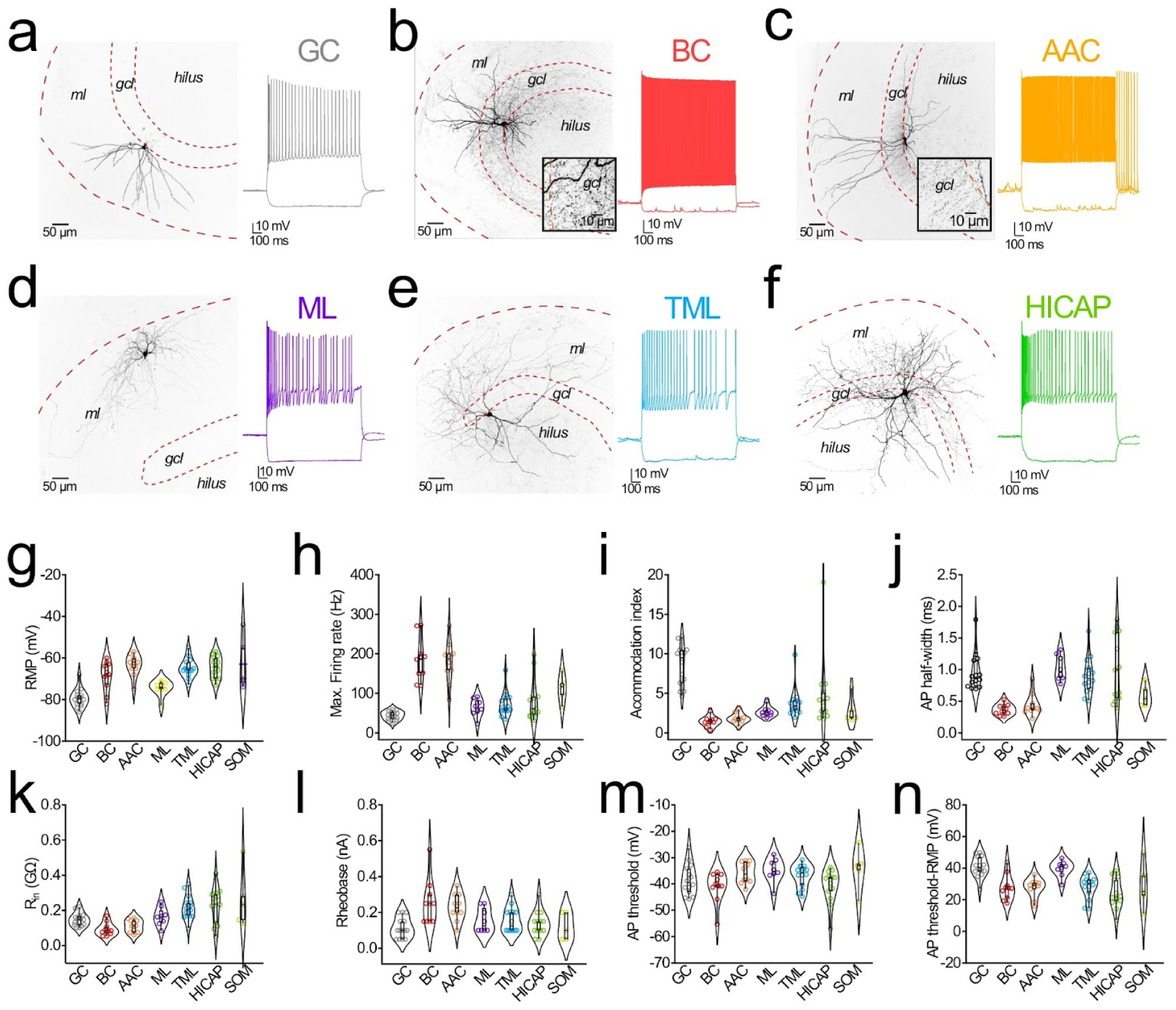
Morphological and physiological properties of DG cell types. **(a-f)** Representative maximum intensity projections of confocal images from a biocytin-labeled granule (GC, **a**), basket (BC, **b**), axo-axonic (AAC, **c**), molecular layer (ML, **d**), total molecular layer (TML, **e**), and hilar-commissural PP-associated cell (HICAP, **f**) with their respective firing patterns. **(g-n)** Physiological properties of recorded DG cell types: (**g**) resting membrane potential (RMP), (**h**) maximum firing rate, (**i**) accommodation ndex, (**j**) action potential half-width, (**k**) input resistance (R_in_), (**l**) rheobase, (**m**) AP threshold, and (**n**) AP threshold - RMP in mV. Notice that RMP, Maximum Firing Rate, and Accommodation Index allowed a reliable differentiation between GCs and INs, as well as between FS-INs (BC and AACs) and RS-INs (TML and HICAPs). Data are presented as violin plots showing the kernel density estimation of the distribution; horizontal lines ndicate the median, and whiskers represent the interquartile range. Individual data points (circles) represent single cells. GC (n=14), BC (n=10), AAC (n=10), ML (n=8), TML (n=14), HICAP (n=13), SOM (n=5).

**Table 1:** Physiological properties of recorded GCs and identified interneurons. Values are given as mean ± standard error of the mean.

| Cell type (n) | RMP (mV) | Max. Firing rate (Hz) | Accommodation index | AP Half-width (ms) | $R_{in}$ (MΩ) | Rheobase (pA) | AP Threshold (mV) | Threshold -RMP (mV) |
| --- | --- | --- | --- | --- | --- | --- | --- | --- |
| GC (14) | -79.3 ± 1.2 | 44 ± 3 | 8.6 ± 0.7 | 0.9 ± 0.08 | 161 ± 14 | 114 ± 15 | -38.5 ± 1.5 | 41 ± 2.6 |
| BC (10) | -68.8 ± 2.3 | 186 ± 17 | 1.4 ± 0.2 | 0.36 ± 0.03 | 142 ± 19 | 255 ± 40 | -41 ± 1.8 | 27.6 ± 2.5 |
| AAC (10) | -63.4 ± 1.7 | 179 ± 16 | 1.8 ± 0.2 | 0.4 ± 0.05 | 162 ± 12 | 230 ± 23 | -35.8 ± 1.3 | 27.6 ± 1.9 |
| ML (8) | -74.5 ± 1.3 | 64 ± 8 | 2.7 ± 0.2 | 1 ± 0.07 | 250 ± 42 | 156 ± 23 | -34.8 ± 1.6 | 39.6 ± 1.8 |
| TML (14) | -64.8 ± 1.2 | 72 ± 8 | 3.8 ± 0.5 | 0.8 ± 0.07 | 246 ± 25 | 150 ± 17 | -36.8 ± 1.1 | 27.8 ± 3 |
| HICAP (13) | -64.5 ± 1.5 | 84 ± 14 | 4.9 ± 1.2 | 1 ± 0.1 | 244 ± 30 | 138 ± 18 | -40.6 ± 1.8 | 23.8 ± 2.6 |
| SOM (5) | -63 ± 5.8 | 112 ± 15 | 2.8 ± 0.7 | 0.5 ± 0.08 | 696 ± 427 | 120 ± 34 | -38.5 ± 1.5 | 28.7 ± 6.5 |

### LEC inputs differentially recruit the different types of DG INs

To understand the mechanisms of maintained sparsity upon LEC stimulation, we performed whole-cell patch-clamp recordings of DG INs while stimulating the outer perforant path. Similar to the Ca^2+^ imaging experiments (**Fig. 1b**), only maximal LPP stimulation (100 V) was able to recruit some GCs (7 out of 12 cells, **Fig. 3a),** which fired APs only sparsely (2.4 ± 0.7 APs per trial, 5 pulses at 30 Hz, **Fig. 3b**), even when cells were held at −70 mV, ∼10 mV more depolarized than their *in vitro* RMP (**Fig. 2g**) and close to their resting potential measured *in vivo* (Pernía-Andrade and Jonas, 2014). This observation is consistent with the finding that several APs are required to induce substantial Ca²⁺ influx in GCs (Stocca, Schmidt-Hieber and Bischofberger, 2008).

**Figure 3:**
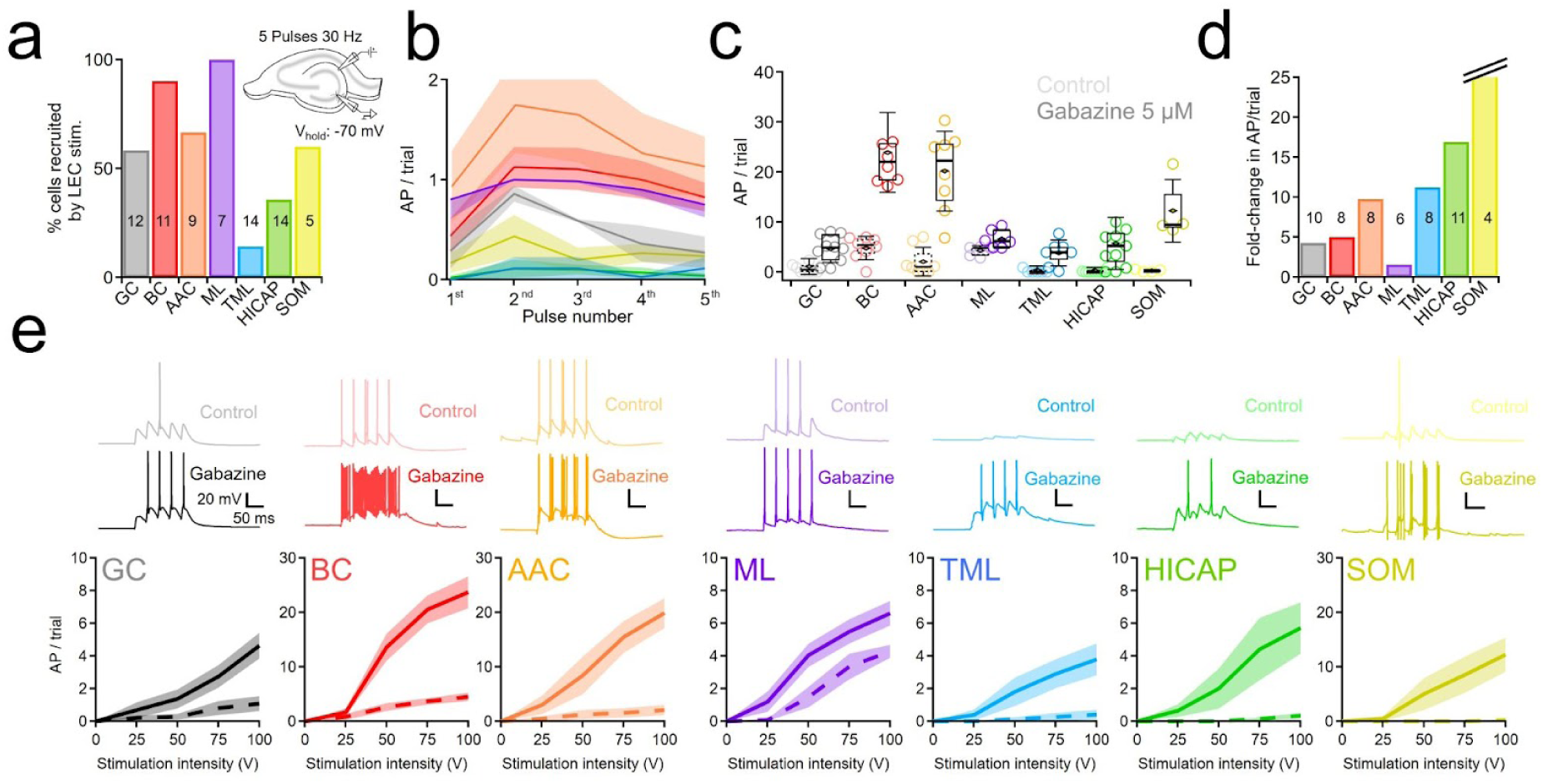
GABA_A_R-mediated inhibition modulates differential LEC-mediated recruitment of the DG circuit. **(a)** Percentage of cells firing at least one action potential (AP) after gamma-modulated stimulation of LEC inputs n control conditions. **(b)** Mean number of AP per trial discharged at every pulse of a 30 Hz stimulation train. **(c)** Box plot showing the number of AP per trial discharged in control conditions (*left boxes*) and after perfusion of gabazine (*right boxes*) per cell type. **(d)** Bar plot showing the average fold-change in AP/trial for every cell type. **(e)** The upper panels show representative traces of membrane potential responses to electrical stimulation in control conditions (light traces) and after perfusion with gabazine (dark traces). Bottom panels show mean AP per trial at the four stimulation intensities tested in control conditions (dashed line) and after GABA_A_R block (solid ines) for the respective cell type. Shades represent the standard error of the mean. Numbers on bars represent the number of recorded cells.

Next, we analyzed which INs were responsible for the sparse GC response following LEC activation by evaluating their recruitability during LPP stimulation. FS-INs were readily recruited after LPP stimulation, with a moderate difference between the two FS-IN subtypes: BCs fired more reliably than AACs (10 out of 11 and 6 out of 9 cells recruited with 5.7 ± 1.2 and 5.5 ± 1.9 APs/trial, respectively, **Fig. 3a, b**). Similarly, ML cells were efficiently recruited by LEC inputs (7 out of 7 cells recruited, with 4.5 ± 0.3 APs/trial) as well as a significant proportion of the tested SOMIs (3 out of 5 cells recruited, with 1.3 ± 0.8 APs/trial). In stark contrast, most TML and HICAP cells were not recruited or fired only a few APs, even at higher stimulation intensities (2 out of 14 and 5 out of 14 cells recruited, firing 1.45 ± 0.5 and 1.16 ± 0.25 APs/trial, respectively), suggesting that LEC inputs differentially engage GABAergic INs in the DG, with FS-INs and ML cells being the most actively driven. AP firing dynamics varied across neuronal types, with FS-INs, ML cells, and GCs discharging during the first pulse of the stimulation train, whereas TML HICAP and SOMI fired only after the second pulse (**Fig. 3b**). Following this initial increase in AP discharge, facilitation either remained stable or slightly declined in all tested DG cell types (**Fig. S1**).

How does GABAergic inhibition control the discharge of the different cells in the DG? To address this question, we tested the effect of blocking GABA_A_Rs on the recruitment of the different DG neurons following LPP stimulation. As expected from the effects observed during Ca²⁺ imaging experiments (**Fig. 1b**), the proportion of recruited GCs as well as their firing rate strongly increased during gabazine application (from 5 out of 10 GCs firing 1 ± 0.5 APs per trial in control conditions to 10/10 cells firing 4.7 ± 0.7 APs per trial in gabazine, maximum stimulation intensity, **Figs. 3c-e**). Similarly, FS-INs substantially increased their firing when GABA_A_R-mediated inhibition was blocked (BCs: 7 out of 8 BCs firing 4.5 ± 0.8 APs/trial, to 8 out of 8 cells firing 24 ± 2.9 APs/trial; AACs: 5 out of 8 AACs firing 2.1 ± 1 APs/trial to 8/8 cells firing 21.5 ± 3.4 APs/trial, for control and gabazine conditions, respectively, **Figs. 3c-e**). FS-INs depolarized to such an extent that depolarization block was frequently observed at higher intensities. MLs displayed only a moderate increase in firing rate after GABA_A_Rs were blocked (ML: from 4.3 ± 0.4 to 6.6 ± 0.8 APs/trial). Remarkably, under these conditions, recruitment of RS-INs was frequently achieved, with a ∼10-fold increase in firing rate (TML: from 2 out of 8 neurons firing 0.4 ± 0.3 APs per trial to 8/8 cells firing 3.8 ± 0.9 APs/trial; HICAP: From 4 out of 11 HICAPs firing 0.3 ± 0.2 APs/trial to 9/11 cells firing 5.7 ± 1.6 APs/trial in control and gabazine conditions, respectively, **Figs. 3c-e**). SOMIs also drastically increased their firing rate upon gabazine application (SOM: From 2 out of 4 SOMIs firing 0.2 ± 0.1 APs/trial to 4/4 cells firing 12.3 ± 3.1 APs/trial, **Fig. 3c-e**). In summary, all investigated cell types increased their discharge rate after GABA_A_R-mediated inhibition was eliminated. The most significant effects were observed in RS-INs and SOM-INs, followed by FS-INs and GCs, while ML layer cells were only slightly affected. Moreover, blocking GABA_A_Rs differentially impacted the dynamics of LEC-induced firing, enhancing facilitation in GCs while reducing it in FS-INs (**Fig. 3b**, **S1**), highlighting that PP-recruited inhibition exhibits distinct activity patterns depending on the specific target cell.

### Differential excitatory drive from LEC into DG INs

What physiological and synaptic properties underlie the differential recruitment of INs by LEC inputs? Neither the resting membrane potential (RMP) nor the AP threshold showed significant differences among the investigated IN types (**Fig. 2g, m**; **Table 1**). The voltage difference between RMP and AP threshold was significantly larger only when comparing GCs to all INs and for ML compared to HICAP cells (**Fig. 2n**; **Table 1**). Input resistance, a parameter highly descriptive of neuronal excitability, was notably lower in FS-INs. At the same time, the rheobase tended to be higher compared to RS-INs (**Fig. 2l, i**; **Table 1**), indicating that intrinsic IN properties might not account for the higher discharge probability observed in FS-INs after LPP stimulation. Therefore, we hypothesized that selective recruitment of INs by the LEC might rely on differences in the properties of synaptic inputs innervating these INs. To test this, we recorded excitatory postsynaptic currents (EPSCs) evoked by LPP stimulation in control conditions and during gabazine perfusion. In control conditions, the amplitude of LEC-mediated EPSCs across the different DG cell types was highly variable (**Fig. 4a**) and generally correlated with the discharge rate observed in these conditions (**Fig. 3c**). BC, GC, AAC, and ML-INs showed the largest EPSC amplitudes (sum of all EPSPs evoked at 100 V intensity: 6 ± 0.9, 3.6 ± 0.3, 3.1 ± 1.4, and 3.7 ± 0.8 nA; n = 6, 10, 6, and 5 cells, respectively, **Fig. 4a**). Consistently with their sparse recruitment, TML and HICAP cells had significantly smaller EPSCs (0.4 ± 0.06 and 0.9 ± 0.2 pA, n = 9 and 7 cells, respectively). This suggests that TML and HICAP cells receive weaker or fewer inputs from the LEC compared to GCs, FS-INs, and ML cells. LEC inputs showed facilitation across all recorded cell types, with TML cells showing a slightly higher, more consistent facilitation ratio (**Figs. 4b-c**). ML INs and GCs showed the lowest facilitation index, which progressively declined with each pulse. Therefore, short-term synaptic dynamics of LEC inputs onto DG neurons cannot explain the differential recruitment of these cells.

**Figure 4:**
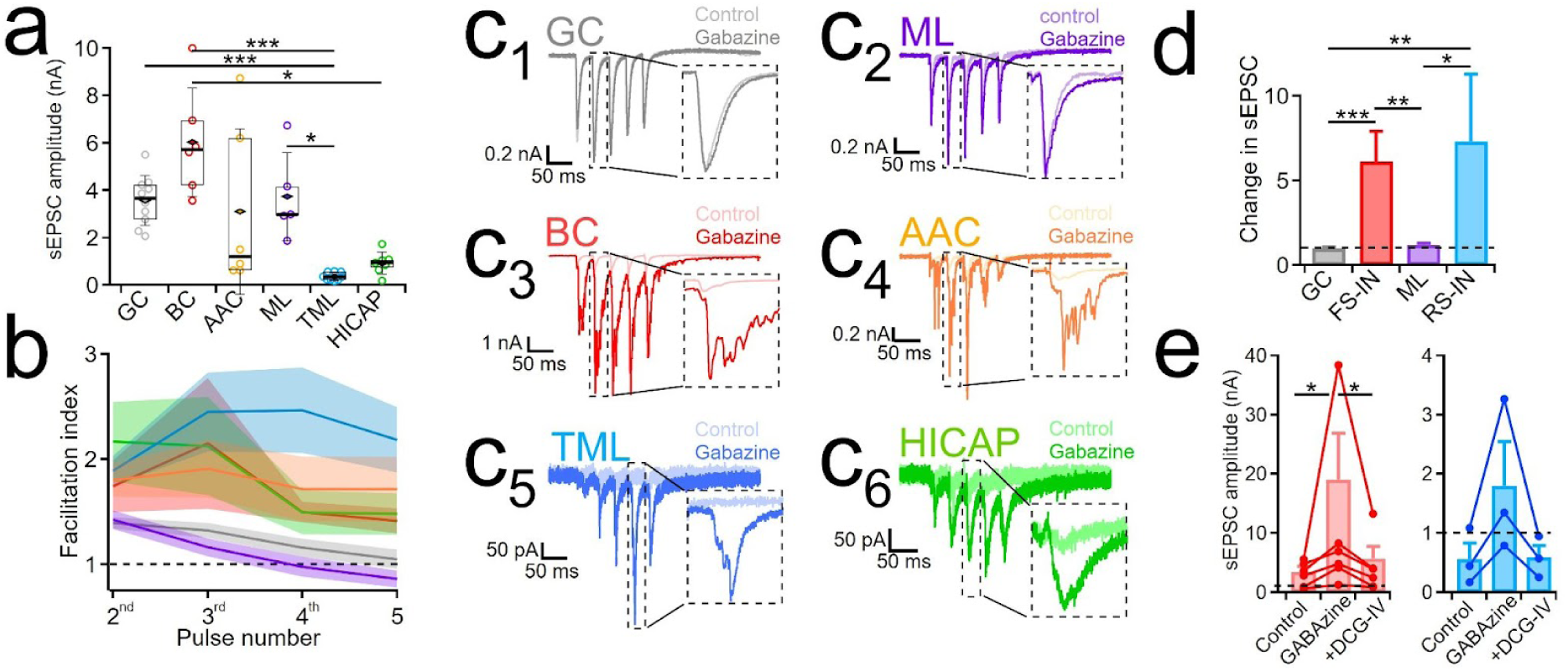
Different LEC feedforward and GC feedback inputs control the differential recruitment of DG neurons. **(a)** Summed amplitude of EPSCs evoked by LPP electrical stimulation at maximal intensity. **(b)** Summary plot showing the facilitation index of summed EPSCs for GCs, BCs, AACs, MLs, TMLs, and HICAPs (p = 1.20 × 10⁻⁵, Kruskal-Wallis test with post-hoc Dunn’s test with Holm corrections, n = 10, 6, 6, 9, and 7, respectively). **(c)** Representative traces of EPSCs evoked by 30 Hz stimulation of the LPP in granule cells (GC), molecular layer cells (ML), basket cells (BC), axo-axonic cells (AAC), total molecular layer cells (TML), and hilar-commissural PP-associated cells (HICAP) in control conditions (light traces) and after perfusion of gabazine (5 µM, dark traces). **(d)** Bar plot showing the mean change in summed EPSC amplitude between control conditions and after gabazine perfusion for GCs, FS-INs, ML-INs, and RS-INs (p = 0.00019, Kruskal-Wallis test with post hoc Dunn comparisons with Holm correction: n = 8, 9, 4, and 11, respectively). **(e)** Bar plot showing the summed mean EPSC amplitude in control conditions after gabazine perfusion and after perfusion with gabazine and DCG-IV for FS-INs (left, p = 0.0036, Friedman test with post hoc Wilcoxon signed-rank tests with Holm correction: n = 6) and for RS-INs (n = 3)., *p < 0.05, **p < 0.01, ***p < 0.001.

Interestingly, although GABA_A_R block induced a significant increase in discharge rate for all DG cell types tested (**Fig. 3**), this manipulation increased EPSC amplitude in FS-INs (from 3.2 ± 0.7 to 16 ± 5.3 nA, p = 0.0039, Wilcoxon signed rank test, n = 8, **Figs. 4c-d**) and RS-INs (from 0.5 ± 0.03 nA to 6.2 ± 1.2 nA, p = 0.0001, Wilcoxon signed rank test, n = 11), but not in GCs or ML-INs (p = 0.5 and 0.8, n = 8 and 4, respectively, **Figs. 4c-d**). While GCs and MLs have only apical dendrites located in the molecular layer, FS- and RS-INs have dendrites located both in the molecular layer and in the hilus (**Figs. 2, 5**), enabling them to receive feedback excitation from GC and mossy cells, which could provide additional excitatory drive during LEC input. Examination of EPSCs after GABA_A_Rs were blocked revealed that they closely resembled mossy-fiber-mediated signaling from GCs to INs: EPSCs were larger, more delayed, and less synchronous (Sambandan et al., 2010) than EPSCs under control conditions (**Fig. 4_c3-c6_**). This suggests that, consistent with **Fig. 1**, increased recruitment during GABA_A_Rs blockade reflects disinhibition—reduced feedback inhibition with a consequent rise in feedback excitation from GCs and mossy cells onto INs. To confirm this, we combined the blockade of GABA_A_Rs with a drug-elicited reduction of glutamate release from GC mossy fibers by adding the specific mGluR type-II agonist DCG-IV (Sambandan *et al*., 2010). This manipulation counteracted the effect of gabazine both in EPSC amplitude (**Fig. 4e**) and in AP firing for FS-INs and RS-INs (FS-INs: Control = 2.34 ± 1.2; gabazine = 19.8 ± 3.5; gabazine + DCG-IV = 12.5 ± 3.4 APs/trial. RS-INs: Control = 0.2 ± 0.1; gabazine = 4 ± 0.8; gabazine + DCG-IV = 2 ± 0.5 APs/trial). Therefore, under vigorous LEC activity, feedback excitation from GCs is a critical excitatory drive that modulates the recruitment of DG INs.

**Figure 5.**
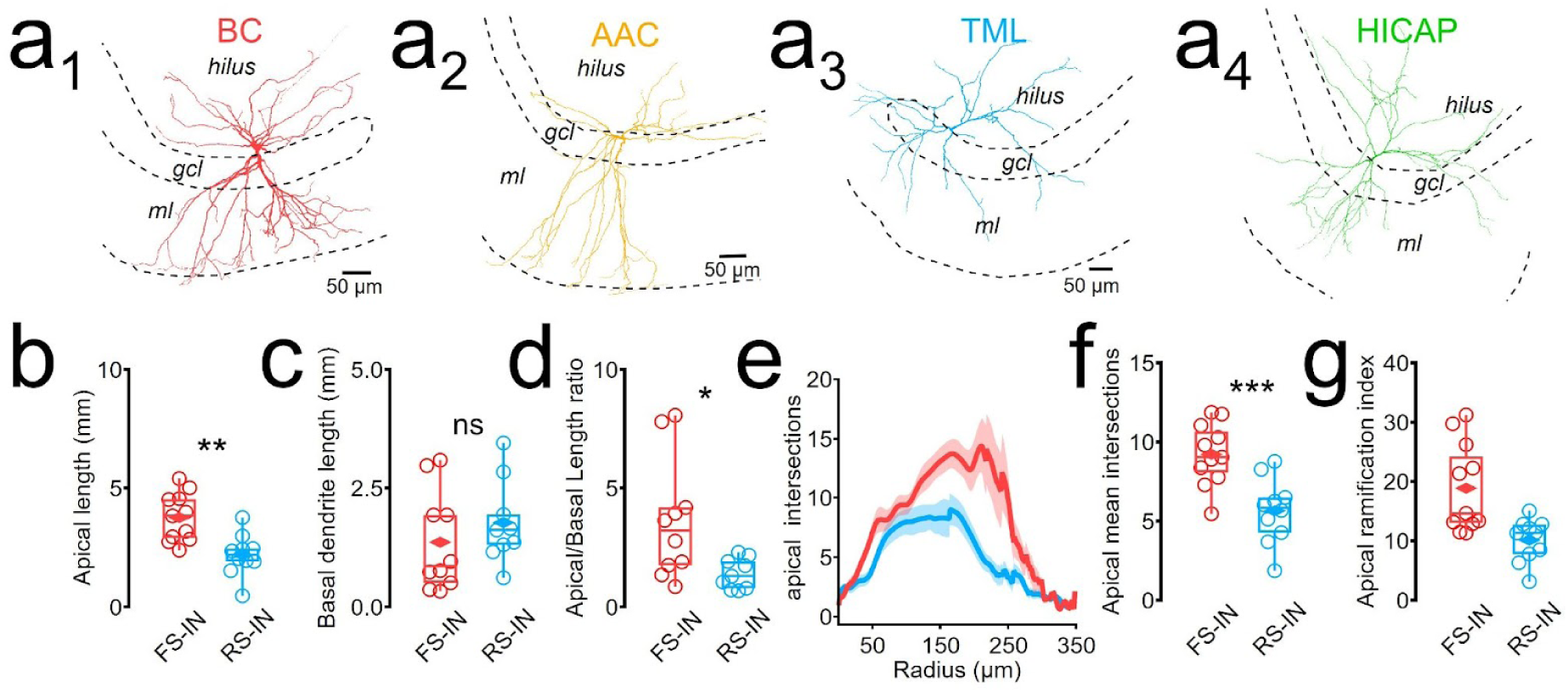
Morphological analysis of fast-spiking and regular-spiking interneurons. **(a)** Maximum projections of reconstructed dendritic processes from representative BC, AAC, TML, and HICAP cells. Notice that dendrites from TML and HICAP cells do not reach the end of the molecular layer. (**b**) Total dendritic length of apical dendrites of FS-INs vs RS-INs (p = 0.001, two-tailed unpaired t-test, 11 and 10 reconstructions, respectively). (**c**) Basal dendrite length of FS-INs vs RS-INs (p = 0.36, two-tailed unpaired t-test, 10 and 9 reconstructions, respectively). (**d**) Apical/basal length ratio for FS vs RS (n=9) interneurons (p = 0.04, two-tailed unpaired Wilcoxon rank sum test, 10 and 9 reconstructions, respectively). Sholl analysis (**e**), mean ntersections (**f**), and apical ramification index (**g**) of FS (n=11) and RS-IN (n=9) apical dendrites. ns = not significant, *p < 0.05, **p < 0.01, ***p < 0.001.

**Figure 6:**
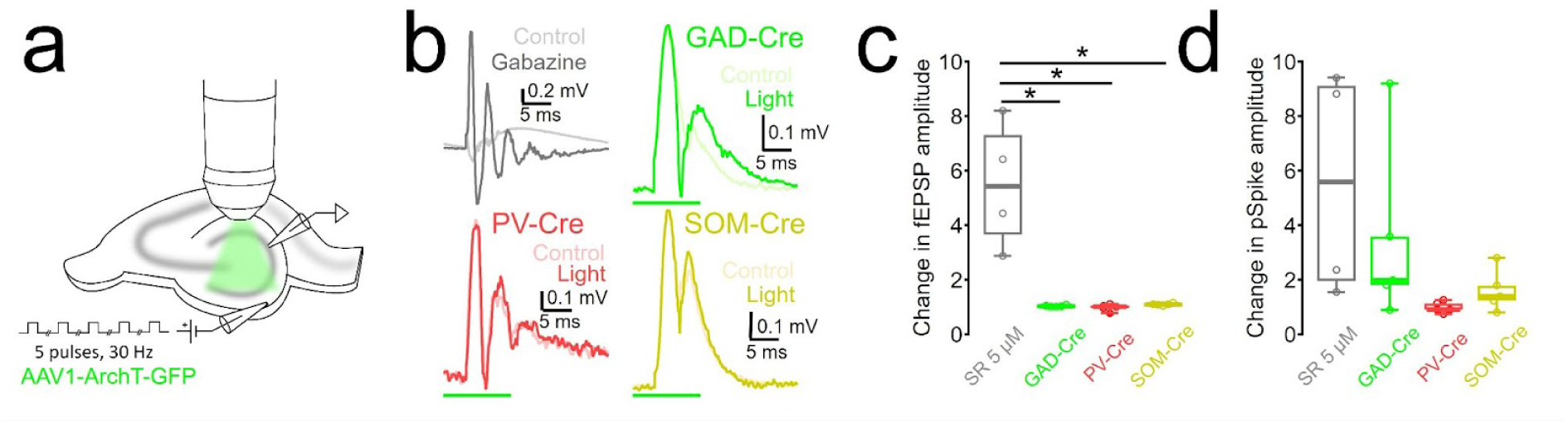
Distinct contribution of IN types in control of DG activity during LEC stimulation. **(a)** Schematic representation of the experiment. Extracellular stimulation (5 pulses at 30 Hz) was applied while recording the extracellular field excitatory postsynaptic potential (fEPSP) in the granule cell layer. Simultaneously, archaerhodopsin (ArchT), virally expressed either in GAD-, PV-, or SOM-Cre mice, was stimulated by wide-field green light to inhibit the target IN population. **(b)** Representative fEPSP traces recorded in the granule cell layer in acute slices after stimulation of the outer molecular layer in control conditions and after perfusion of gabazine (*top eft*), or after ArchT-mediated inhibition of GAD2- (*top right*), PV- (*bottom left*), or SOM-expressing (*bottom right*) INs. **(c-d)** Summary box plots showing the effect of applying gabazine (grey, n = 4) and the impact of optogenetic nhibition in GAD- (green, n = 5), PV- (red, n = 5), or SOM- (yellow, n = 5) expressing INs on the amplitude of the fEPSP (**c**) and on the amplitude of the population spike (**d**). Kruskal-Wallis with Games-Howell post hoc tests. * p < 0.05.

To understand why LEC stimulation preferentially recruits FS-INs compared to RS-INs, we reconstructed the dendrites of biocytin-filled FS- and RS-INs and measured their morphological properties (**Fig. 5a**). Apical dendrites of FS-INs were significantly longer than those of RS-INs (3.3 ± 0.3 vs 2.2 ± 0.3 mm, p = 0.001, **Fig. 5b**), while their basal dendrites did not vary from each other (1.4 ± 0.3 vs 1.8 ± 0.3 mm, **Fig. 5c**), resulting in a significantly larger apical/basal dendritic length ratio for FS-INs (5.2 ± 1.5 vs 1.7 ± 0.3 mm, **Fig. 5d**). Sholl analysis of apical dendritic trees also revealed that the apical dendrites of FS-INs extended further into the molecular layer than RS-INs (11.2 ± 2.4 vs. 4.3 ± 0.8 mean intersections 200–250 µm from the soma, p = 0.01, n = 11 and 9, two-tailed unpaired Wilcoxon Rank Sum test, **Fig. 5a** and **e**). Moreover, FS-INs exhibited a more complex apical dendritic architecture as indicated by the mean number of intersections across the entire apical arbor (9.2 ± 0.6 vs. 5.6 ± 0.7 intersections, p = 0.0006, n = 11 and 9, two-tailed unpaired t-test; **Fig. 5f**) and the Sholl ramification index (18.9 ± 2.2 vs. 10.1 ± 1.1, p = 0.003, n = 11 and 9, two-tailed unpaired Wilcoxon rank sum test; **Fig. 5g**). In summary, FS-INs have more complex apical dendrites that extend further into the molecular layer, which makes them ideally positioned to receive LEC synaptic contacts. RS-INs, on the other hand, extend their dendrites in more proximal regions of the molecular layer and in the hilus, where they may preferentially form synapses with MEC, mossy cell, or feedback mossy fiber axonal projections.

### Differential impact of IN types on GC recruitment by LEC inputs

To test directly which INs are engaged in controlling DG population activity during strong LEC drive, we used an optogenetic approach combined with recordings of evoked local field potentials (LFPs) in response to 30 Hz stimulation of LEC inputs (Lee et al., 2016, **Fig. 6a**). First, we evaluated the effect of GABA_A_R-mediated inhibition on the GC population. As expected, blocking GABA_A_Rs produced an substantial increase in both the amplitude of the field EPSP (fEPSP) and the population spike (pSpike) recorded in the gcl (5.5 ± 1.2 and 5.5 ± 2-fold increase in the amplitude of the fEPSP and pSpike for the 2^nd^ stimulation pulse, p = 0.016 and 0.03, respectively, n = 4, **Figs. 6b-d**), consistent with the impact on GCs Ca^²⁺^-transients during gabazine application (**Fig. 1**). Second, we expressed the light-sensitive proton-pump archaerhodopsin (ArchT) in GAD2-Cre mice to evaluate the impact of inhibiting a large and heterogeneous IN population and in PV- or SOM-Cre mice to test the effect of inhibiting perisomatic and dendritic-targeting INs, which can be recruited during LPP activation (**Fig. 3**). ArchT-mediated inhibition of GAD2-expressing INs produced a moderate change and affected the fEPSP and pSpike amplitude (from 2.5 ± 0.2 to 2.6 ± 0.2 mV and from 0.15 ± 0.05 to 0.3 ± 0.1 mV, p = 0.03 for both, n = 5, **Fig. 6b-d**). Interestingly, inhibiting release from ArchT-expressing FS-INs had no effect on the fEPSP amplitude or the pSpike, (p = 0.5 for both), while disrupting GABA release from SOMIs had a mild but significant effect on the GC population response (from 2.4 ± 0.4 to 2.6 ± 0.04 mV and from 0.14 ± 0.07 to 0.18 ± 0.09 mV, p = 0.0005 and 0.0004 for fEPSP and pSpike, respectively, n = 5, **Fig. 6b-d**). These results suggest that, although both perisoma- and dendrite-targeting inhibitory INs are recruited during LEC stimulation, the latter have a more significant effect on the recruitment of GCs after stimulation of their distal dendrites. Moreover, it also shows that pharmacologically blocking all GABA_A_R receptors has a much larger impact than the transient silencing of a substantial fraction of INs, suggesting cooperative interneuron activity or tonic GABA_A_R activation may have an important role in maintaining sparsity within the DG. Cooperative regulation of GC activity by distinct IN types—via complex disinhibitory circuits—may have contributed to these observations and influenced our optogenetic manipulations.

## Discussion

The sparse activity of GCs is an essential condition for the generation of an orthogonalized representation of incoming entorhinal information and the DG’s pattern separation function (Rolls, 2013). The sparsity of the DG code is likely mediated by the intrinsic properties of adult GCs and the tight inhibitory control provided by different types of GABAergic INs (Hainmueller *et al*., 2024b). Because DG INs are themselves strongly inhibited, the resulting lateral and disinhibitory interactions may modulate the magnitude and dynamics of DG activity. Here, we show that LEC inputs differentially recruit IN types in the DG, thereby contributing to type-specific control of GC activity. Gamma-frequency range LEC stimulation readily recruited FS-INs (BCs, AACs), consistent with their role in providing soma-near feedforward inhibition. By rapidly and powerfully inhibiting GC somata and proximal dendrites, BCs and AACs likely prevent most GCs from reaching spike threshold during initial input volleys. Concurrently, ML-INs, which are best positioned to enable feed-forward inhibition, were also reliably activated by incoming PP activity, imposing an additional control on GC dendritic integration. This feedforward recruitment of both perisomatic and dendritic inhibition explains why, in control conditions, even strong LEC-driven synaptic input translates into minimal GC firing as proximal inhibition raises the spiking threshold of GCs, while distal inhibition dampens excitatory synaptic integration in the dendrites. In contrast, RS-INs and SOMIs showed little or no spiking in response to the same LEC stimulation. These cells were recruited only when GABA_A_Rs were blocked or at very high stimulus intensities, suggesting they primarily participate in feedback inhibition. By engaging in the network only when GCs are firing, they provide a secondary source of inhibition that may help restrain GC activity if the initial gate is insufficient—protecting the DG against overexcitation and pathological states such as seizures or excitotoxicity (Dengler *et al*., 2017). The demonstrated differential recruitment of IN classes by the LEC is unlikely to be supported by intrinsic physiological differences (**Fig. 2 and Table 1**). Instead, it appears to reflect their distinct morphological and synaptic features, such as the larger coverage of the oml by the apical dendrites of FS-INs compared to RS-INs (**Fig. 5**) and the lower initial discharge probability of RS-INs (**Fig. 4**). Previous studies have shown that a similar preferential recruitment of FS-INs over RS-INs occurs upon MEC stimulation (Liu, Cheng and Lien, 2014; Hsu et al., 2016), supporting the idea that the feedforward versus feedback functional distinction between these IN types is preserved regardless of the specific source of cortical input, although the mechanisms involved might be different. FS-INs have low input resistance and rapid membrane time constants, requiring strong synaptic input to discharge, but they respond with minimal delay and high-frequency outputs (Bartos *et al*., 2002; Jonas *et al*., 2004). These properties make them ideal for rapid feedforward inhibition, controlling spike timing and the synchronization of neuronal networks at gamma frequency (Bartos *et al*., 2002). In contrast, many dendrite-targeting interneurons (e.g., SOMI: hilar perforant path-associated cells or neurogliaform cells) are regular-spiking or adapting, with higher input resistance and longer integration windows (Armstrong *et al*., 2011; Yuan *et al*., 2017). They will reach firing threshold more readily with incremental or sustained input, and their inhibition can be more prolonged. This allows them to integrate slower or repetitive activity and provide inhibition over a longer timescale (e.g., across theta cycles or during bursts), complementing the phasic, rapid inhibition provided by FS-INs. In a spiking neural network model, the division in soma-controlling and dendrite-controlling INs emerged as an optimal computational solution for providing independent, compartment-specific feedback inhibition to principal cells (Keijser and Sprekeler, 2022). Thus, compartment-specific recruitment may tune IN subtypes to distinct spatiotemporal features of network activity, collectively covering a wide range of excitatory drive. Our results also highlight that DG INs not only inhibit GCs but also each other (Bartos *et al*., 2002), forming complex disinhibitory motifs. The pronounced increase in IN spiking across all types when GABA_A_Rs were blocked suggests that there are significant inhibitory interactions between INs. Blocking GABA_A_Rs produced broad coactivation of INs and sharply increased spiking across all IN types, indicating potent mutual inhibition under normal conditions. These findings are in line with prior studies demonstrating that SOMIs and PVIs can bidirectionally and powerfully inhibit each other (Savanthrapadian *et al*., 2014; Yuan *et al*., 2017; Elgueta and Bartos, 2019), which could create a regulatory loop: When LEC activity is low, inhibition is mainly provided by FS-INs, which concomitantly inhibit SOMIs. However, when network activity rises, SOMIs are strongly recruited. In turn, they can transiently inhibit PV-INs, thereby easing the suppression of GC somas while keeping local dendritic activation to a minimum and shifting the locus of inhibition from proximal to distal (Pouille and Scanziani, 2004). Therefore, IN-IN inhibition allows the circuit to dynamically reconfigure via disinhibition, tightening or loosening the gate on GC activity depending on the context, rather than acting as a static blanket of inhibition (Karnani *et al*., 2014). Our results substantiate this interpretation by showing that pharmacological removal of inhibition unmasked a latent excitatory drive in the network, evident in both principal cells and INs. In physiological conditions, more subtle shifts—for instance, cholinergic modulation or novelty exposure (Hainmueller *et al*., 2024b)—might similarly tip the balance between inhibition and disinhibition, thereby tuning how permissive the DG is to GC discharge.

Importantly, feedforward and feedback inhibition in the DG operate in concert with disinhibitory motifs. Our analysis (**Fig. 6**) indicated that blocking GABA_A_Rs preferentially enhanced delayed, polysynaptic EPSCs onto INs—presumably arising from GC or mossy cell feedback excitation—rather than immediate monosynaptic EPSCs from PP inputs. Thus, gabazine increased interneuron recruitment primarily by disinhibiting GCs, which then drove INs via feedback excitation. We believe that under behavioral conditions, similar mechanisms might occur in a more subtle, nuanced manner. As selected, sparse populations of GCs overcome inhibition, they will recruit RS-INs and SOM-INs, thereby effectively shifting the locus of inhibition towards GC dendrites. This self-regulating feedback loop may prevent further dendritic plasticity in cells that are already part of a DG engram (Schmidt-Hieber *et al*., 2004; Lopez-Rojas *et al*., 2016). Disinhibitory connections might transiently delay this feedback or target it to specific compartments by selecting specific IN types, ensuring that the inhibition arrives in a controlled fashion and allowing a brief window for some GCs to transmit information to downstream CA3 before inhibition is fully engaged. Such timing-dependent interactions likely prevent saturation during bursts and preserve sparsity across a wide range of input strengths, thereby enabling the DG to act as the gate of the trisynaptic hippocampal loop while simultaneously supporting pattern separation.

## Material and Methods

### Slice preparation and electrophysiology

Transverse acute hippocampal slices (300–350 µm) were obtained from 3-10 week old mice from either sex in accordance with national and institutional legislation (license X-13/03S, X-16/306S, and G-15/106 approved by the Regierungspräsidium Freiburg). Slice preparation and whole-cell patch-clamp recordings were performed as previously described (Elgueta et al., 2015) using cold sucrose-or NMDG-based ACSF after intracardial perfusion (Ting et al., 2014). All recordings were performed in ACSF containing (in mM): 125 NaCl, 25 NaHCO3, 2.5 KCl, 1.25 NaH2PO4, 25 d-glucose, 2 CaCl2, and 1 MgCl2 (oxygenated with 95% O2/5% CO2, ∼31-33°C). Patch pipettes were pulled from borosilicate glass tubing (Flaming-Brown P-97 puller, Sutter Instruments) and had 2.7–6.0 MOhm resistance after filling with an intracellular solution containing (in mM): 120 K-gluconate, 20 KCl, 10 HEPES, 2 MgCl_2_, 2 Na2 ATP, 10 EGTA, and 0.2% biocytin (pH 7.2, 290-310 mOsm). Neurons in the dentate gyrus (DG) were approached under visual control using infrared DIC microscopy (Zeiss Axio Examiner). Whole-cell current- and voltage-clamp recordings were obtained using a Multiclamp 700B amplifier (Molecular Devices), filtered at 5 kHz, and digitized at 20-40 kHz (Power1401-2 interface; Cambridge Electronic Design). Series resistance (4–20 MΩ) was compensated in current-clamp mode but not in voltage-clamp mode (holding potential: −70 mV). Intrinsic properties of recorded cells were measured right after obtaining the whole-cell configuration. Stimulus generation and data acquisition were performed with a custom-made program (FPulse) written in Igor Pro 6.22 (Wavemetrics). Gamma-modulated lateral entorhinal cortex inputs into the DG were emulated using a glass tubing stimulation pipette placed in the outer-molecular layer using trains of 5 pulses at 30Hz. To block GABAergic IN activity, slices were perfused with Gabazine (SR-95531, 5 µm).

### Calcium imaging

Recombinant adeno-associated viruses (rAAVs) encoding GCaMP5 (AAV1.hSynap.GCaMP5G, 5 x 10^11^ GC/ml) were stereotactically injected into the ventral DG (relative to Bregma: y, 2.55 mm; x, 2.9 mm; z, 2.3–2.9 mm, 800 nL) of adult C57Bl6/J mice. Slices were prepared >14 d after injection and recorded using a galvanometric-scanning 2-photon microscope (Femto-Alba, Femtonics). GCamp5 fluorescence was stimulated at a wavelength of 850 nm using a Chameleon II 2-photon laser (Coherent). The change in signals across the granule cell layer was recorded after stimulating the outer molecular layer with gamma-modulated signals. ROIs encompassing individual granule cells were manually selected, and dF/F was calculated after background subtraction.

### Optogenetics and field potentials

GAD2-Cre (Gad2<tm2(cre)Zjh>/J), PV-Cre (Pvalb<tm1(cre)Arbr>), and SOM-Cre (see https://www.jneurosci.org/content/jneuro/34/24/8197.full.pdf) mice were injected in ventral DG with recombinant rAAVs encoding the light-sensitive proton-pump Archaerhodopsin and GFP (AAV1-CAG-archT-GFP-WPRE-SV40, 8 x 10^11^ GC), as previously described. ArchT was stimulated using an LED light source (CoolLed, 542 nm, 20 ms) filling the back aperture of the objective and triggered by the recording software. Light inhibition of INs with ArchT was evoked 10 ms before each pulse of 5 LPP stimulation events (30 Hz). DG activity was recorded by placing a glass electrode in the granule cell layer. Recorded signals during stimulation show the population GC depolarization as field EPSPs, as well as synchronous GC action potentials as population spikes superimposed over the field EPSPs.

### Neuronal staining and confocal imaging

After in vitro experiments, slices were fixed overnight in 4% paraformaldehyde and subsequently washed in 0.1 M phosphate-buffered (24 h) and 0.025 M phosphate-buffered solution (PBS, pH 7.3) at room temperature. Confocal images of neurons filled with biocytin were taken using a laser scanning confocal microscope (LSM-710, Zeiss) with 20x or 40x magnification. The obtained morphological data were used to classify cells based on their relative position in the dentate gyrus and their axonal morphology.

### Data analysis

Physiological data were measured and analyzed using custom-made IgorPro analysis routines (Wavemetrics). Calcium imaging data were analyzed using custom-written MATLAB routines and the MES 6 package from Femtonics. In some cases, during block of GABA_A_R-mediated inhibition, LEC stimulation induced such strong excitation that FS-cells fired complex spikes or persistent firing was generated, making it difficult to measure individual APs. In those cases, we used a lower intensity, which generated a measurable firing rate. The population spike was determined from the second stimulation event, as it more consistently showed a prominent signal. The population spike amplitude was measured as the biggest drop from the EPSP baseline, which was identified where the EPSP waveform line crossed the baseline. Significance levels are indicated as p-values. Data are presented as mean ± s.e.m., either as bars or shades.

## Supplemental Material

**Figure S1:**
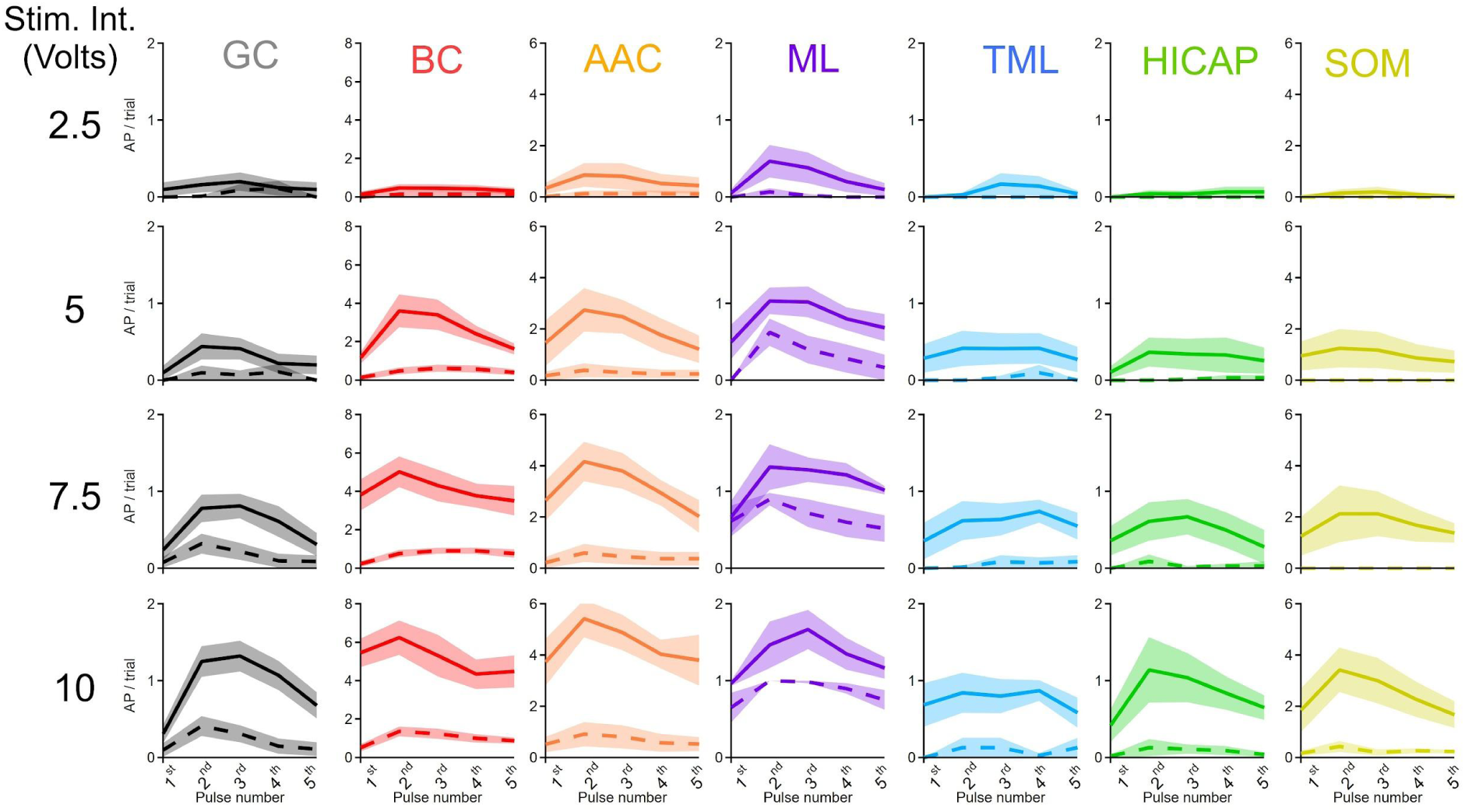
LEC-driven action potential firing dynamics in DG INs and GCs. In control conditions (dashed ines), LEC stimulation at different intensities induces AP discharges in GCs, FSs, and ML-INs. After inhibition was blocked with gabazine (solid lines), all cells fired, although still at different levels (n = 10, 8, 8, 6, 9, 11 and 5 for GC, BC, AAC, ML, TML, HICAP and SOM cells).

## Notes

**Funding statement:** This work was funded by SFB transregio 384 (M.B., C.E.) and the Molecular and Translational Research in Freiburg (MOTI-VATE) Program of the Medical Faculty, University of Freiburg (Else Kröner-Fresenius Foundation; J.K.).

### Competing Interest Statement

The authors have declared no competing interest.

## References

Allegra, M. et al. (2020) ‘Differential Relation between Neuronal and Behavioral Discrimination during Hippocampal Memory Encoding’, Neuron, 108(6), pp. 1103–1112.e6. Available at: 10.1016/j.neuron.2020.09.032.

Armstrong, C. et al. (2011) ‘Neurogliaform cells in the molecular layer of the dentate gyrus as feed-forward γ-aminobutyric acidergic modulators of entorhinal–hippocampal interplay’, Journal of Comparative Neurology, 519(8), pp. 1476–1491. Available at: 10.1002/cne.22577.

Bartos, M. et al. (2001) ‘Rapid Signaling at Inhibitory Synapses in a Dentate Gyrus Interneuron Network’, The Journal of Neuroscience, 21(8), pp. 2687–2698. Available at: 10.1523/JNEUROSCI.21-08-02687.2001.

Bartos, M. et al. (2002) ‘Fast synaptic inhibition promotes synchronized gamma oscillations in hippocampal interneuron networks’, Proceedings of the National Academy of Sciences, 99(20), pp. 13222–13227. Available at: 10.1073/pnas.192233099.

Bartos, M. and Elgueta, C. (2012) ‘Functional characteristics of parvalbumin- and cholecystokinin-expressing basket cells: Parvalbumin- and cholecystokinin-expressing basket cells’, The Journal of Physiology, 590(4), pp. 669–681. Available at: 10.1113/jphysiol.2011.226175.

Bartos, M., Vida, I. and Jonas, P. (2007) ‘Synaptic mechanisms of synchronized gamma oscillations in inhibitory interneuron networks’, Nature Reviews Neuroscience, 8(1), pp. 45–56. Available at: 10.1038/nrn2044.

Cholvin, T., Hainmueller, T. and Bartos, M. (2021) ‘The hippocampus converts dynamic entorhinal inputs into stable spatial maps’, Neuron, 109(19), pp. 3135–3148.e7. Available at: 10.1016/j.neuron.2021.09.019.

Dengler, C.G. et al. (2017) ‘Massively augmented hippocampal dentate granule cell activation accompanies epilepsy development’, Scientific Reports, 7, p. 42090. Available at: 10.1038/srep42090.

Deshmukh, S.S. and Knierim, J.J. (2011) ‘Representation of Non-Spatial and Spatial Information in the Lateral Entorhinal Cortex’, Frontiers in Behavioral Neuroscience, 5. Available at: 10.3389/fnbeh.2011.00069.

Diamantaki, M. et al. (2016) ‘Priming Spatial Activity by Single-Cell Stimulation in the Dentate Gyrus of Freely Moving Rats’, Current Biology, 26(4), pp. 536–541. Available at: 10.1016/j.cub.2015.12.053.

Elgueta, C. and Bartos, M. (2019) ‘Dendritic inhibition differentially regulates excitability of dentate gyrus parvalbumin-expressing interneurons and granule cells’, Nature Communications, 10(1), p. 5561. Available at: 10.1038/s41467-019-13533-3.

Elgueta, C., Köhler, J. and Bartos, M. (2015) ‘Persistent Discharges in Dentate Gyrus Perisoma-Inhibiting Interneurons Require Hyperpolarization-Activated Cyclic Nucleotide-Gated Channel Activation’, The Journal of Neuroscience, 35(10), pp. 4131–4139. Available at: 10.1523/JNEUROSCI.3671-14.2015.

Hainmueller, T. et al. (2024a) ‘Subfield-specific interneuron circuits govern the hippocampal response to novelty in male mice’, Nature Communications, 15(1), p. 714. Available at: 10.1038/s41467-024-44882-3.

Hainmueller, T. et al. (2024b) ‘Subfield-specific interneuron circuits govern the hippocampal response to novelty in male mice’, Nature Communications, 15(1), p. 714. Available at: 10.1038/s41467-024-44882-3.

Hainmueller, T. and Bartos, M. (2018) ‘Parallel emergence of stable and dynamic memory engrams in the hippocampus’, Nature, 558(7709), pp. 292–296. Available at: 10.1038/s41586-018-0191-2.

Hargreaves, E.L. et al. (2005) ‘Major Dissociation Between Medial and Lateral Entorhinal Input to Dorsal Hippocampus’, Science, 308(5729), pp. 1792–1794. Available at: 10.1126/science.1110449.

Hefft, S. and Jonas, P. (2005) ‘Asynchronous GABA release generates long-lasting inhibition at a hippocampal interneuron–principal neuron synapse’, Nature Neuroscience, 8(10), pp. 1319–1328. Available at: 10.1038/nn1542.

Hosp, J.A. et al. (2014) ‘Morpho-physiological criteria divide dentate gyrus interneurons into classes’, Hippocampus, 24(2), pp. 189–203. Available at: 10.1002/hipo.22214.

Jonas, P. et al. (2004) ‘Interneuron Diversity series: Fast in, fast out – temporal and spatial signal processing in hippocampal interneurons’, Trends in Neurosciences, 27(1), pp. 30–40. Available at: 10.1016/j.tins.2003.10.010.

Karnani, M.M., Agetsuma, M. and Yuste, R. (2014) ‘A blanket of inhibition: functional inferences from dense inhibitory connectivity’, Current Opinion in Neurobiology, 26, pp. 96–102. Available at: 10.1016/j.conb.2013.12.015.

Krueppel, R., Remy, S. and Beck, H. (2011) ‘Dendritic Integration in Hippocampal Dentate Granule Cells’, Neuron, 71(3), pp. 512–528. Available at: 10.1016/j.neuron.2011.05.043.

Leutgeb, J.K. et al. (2007) ‘Pattern Separation in the Dentate Gyrus and CA3 of the Hippocampus’, Science, 315(5814), pp. 961–966. Available at: 10.1126/science.1135801.

Liu, X. et al. (2012) ‘Optogenetic stimulation of a hippocampal engram activates fear memory recall’, Nature, 484(7394), pp. 381–385. Available at: 10.1038/nature11028.

Lopez-Rojas, J., Heine, M. and Kreutz, M.R. (2016) ‘Plasticity of intrinsic excitability in mature granule cells of the dentate gyrus’, Scientific Reports, 6(1), p. 21615. Available at: 10.1038/srep21615.

McHugh, T.J. et al. (2007) ‘Dentate Gyrus NMDA Receptors Mediate Rapid Pattern Separation in the Hippocampal Network’, Science, 317(5834), pp. 94–99. Available at: 10.1126/science.1140263.

Morales, C. et al. (2021) ‘Dentate Gyrus Somatostatin Cells are Required for Contextual Discrimination During Episodic Memory Encoding’, Cerebral Cortex, 31(2), pp. 1046–1059. Available at: 10.1093/cercor/bhaa273.

Pernía-Andrade, A.J. and Jonas, P. (2014) ‘Theta-Gamma-Modulated Synaptic Currents in Hippocampal Granule Cells In Vivo Define a Mechanism for Network Oscillations’, Neuron, 81(1), pp. 140–152. Available at: 10.1016/j.neuron.2013.09.046.

Pouille, F. and Scanziani, M. (2004) ‘Routing of spike series by dynamic circuits in the hippocampus’, Nature, 429(6993), pp. 717–723. Available at: 10.1038/nature02615.

Rolls, E.T. (2013) ‘The mechanisms for pattern completion and pattern separation in the hippocampus’, Frontiers in Systems Neuroscience, 7. Available at: 10.3389/fnsys.2013.00074.

Sambandan, S. et al. (2010) ‘Associative plasticity at excitatory synapses facilitates recruitment of fast-spiking interneurons in the dentate gyrus’, The Journal of Neuroscience: The Official Journal of the Society for Neuroscience, 30(35), pp. 11826–11837. Available at: 10.1523/JNEUROSCI.2012-10.2010.

Savanthrapadian, S. et al. (2014) ‘Synaptic Properties of SOM- and CCK-Expressing Cells in Dentate Gyrus Interneuron Networks’, Journal of Neuroscience, 34(24), pp. 8197–8209. Available at: 10.1523/JNEUROSCI.5433-13.2014.

Scharfman, H.E. (2016) ‘The enigmatic mossy cell of the dentate gyrus’, Nature Reviews Neuroscience, 17(9), pp. 562–575. Available at: 10.1038/nrn.2016.87.

Schmidt-Hieber, C., Jonas, P. and Bischofberger, J. (2004) ‘Enhanced synaptic plasticity in newly generated granule cells of the adult hippocampus’, 429.

Schmidt-Hieber, C., Jonas, P. and Bischofberger, J. (2007) ‘Subthreshold Dendritic Signal Processing and Coincidence Detection in Dentate Gyrus Granule Cells’, The Journal of Neuroscience, 27(31), pp. 8430–8441. Available at: 10.1523/JNEUROSCI.1787-07.2007.

Senzai, Y. and Buzsáki, G. (2017) ‘Physiological Properties and Behavioral Correlates of Hippocampal Granule Cells and Mossy Cells’, Neuron, 93(3), pp. 691–704.e5. Available at: 10.1016/j.neuron.2016.12.011.

Stefanelli, T. et al. (2016) ‘Hippocampal Somatostatin Interneurons Control the Size of Neuronal Memory Ensembles’, Neuron, 89(5), pp. 1074–1085. Available at: 10.1016/j.neuron.2016.01.024.

Strüber, M. et al. (2017) ‘Distance-dependent inhibition facilitates focality of gamma oscillations in the dentate gyrus’, Nature Communications, 8(1), p. 758. Available at: 10.1038/s41467-017-00936-3.

Strüber, M., Jonas, P. and Bartos, M. (2015) ‘Strength and duration of perisomatic GABAergic inhibition depend on distance between synaptically connected cells’, Proceedings of the National Academy of Sciences, 112(4), pp. 1220–1225. Available at: 10.1073/pnas.1412996112.

Treves, A. and Rolls, E.T. (1994) ‘Computational analysis of the role of the hippocampus in memory’, Hippocampus, 4(3), pp. 374–391. Available at: 10.1002/hipo.450040319.

Wang, C. et al. (2018) ‘Egocentric coding of external items in the lateral entorhinal cortex’, Science, 362(6417), pp. 945–949. Available at: 10.1126/science.aau4940.

Witter, M.P. (2007) ‘The perforant path: projections from the entorhinal cortex to the dentate gyrus’, in H.E. Scharfman (ed.) Progress in Brain Research. Elsevier (The Dentate Gyrus: A Comprehensive Guide to Structure, Function, and Clinical Implications), pp. 43–61. Available at: 10.1016/S0079-6123(07)63003-9.

Yuan, M. et al. (2017) ‘Somatostatin-positive interneurons in the dentate gyrus of mice provide local- and long-range septal synaptic inhibition’, eLife, 6, p. e21105. Available at: 10.7554/eLife.21105.

Zhang, X., Schlögl, A. and Jonas, P. (2020) ‘Selective Routing of Spatial Information Flow from Input to Output in Hippocampal Granule Cells’, Neuron, 107(6), pp. 1212–1225.e7. Available at: 10.1016/j.neuron.2020.07.006.

